# Spontaneous alpha-band amplitude predicts subjective visibility but not discrimination accuracy during high-level perception

**DOI:** 10.1101/2020.07.13.201178

**Authors:** Jason Samaha, Joshua J. LaRocque, Bradley R. Postle

## Abstract

Near-threshold perception is a paradigm case of perceptual reports diverging from reality – perception of an unchanging stimulus can vacillate from undetected to clearly perceived. Among the many factors that predict whether a stimulus will reach awareness, the amplitude of low-frequency brain oscillations - particularly in the alpha frequency band (8-13 Hz) - has emerged as a reliable predictor of trial-to-trial variability in perceptual decisions. Analysis grounded in signal detection theory suggest that strong prestimulus alpha oscillations diminish subjective perception without affecting the accuracy or sensitivity (d’) of perceptual decisions. These results, coupled with recent studies on sensory responses, point to an inhibitory influence of alpha-band amplitude on early visuocortical activity. The findings to date have been based on simple, low-level visual stimuli, which warrant a focus on early visual processing. However, the physiology of alpha in higher-level visual areas is known to be distinct from early visual cortex, with evidence indicating that alpha amplitude in the inferior temporal (IT) cortex is excitatory (rather than inhibitory, as in early visual cortex). Here, we addressed the question of how spontaneous oscillatory amplitude impacts subjective and objective aspects of perception using a high-level perceptual decision task. Human observers completed a near-threshold face/house discrimination task with subjective visibility ratings while electroencephalograms (EEG) were recorded. Using a single-trial multiple regression analysis, we found that spontaneous fluctuations in pre-stimulus alpha-band amplitude were negatively related to visibility ratings but did not predict trial-by-trial accuracy. These results suggest that the inhibitory influence of prestimulus alpha activity in early visual cortex, rather than the excitatory influence of alpha in IT, comes to bias high-level perceptual reports. Our findings provide further evidence that ongoing alpha amplitude dissociates subjective and objective measures of visual perception.

## 1. Introduction

Conscious perception is determined not only by physical stimulus properties, but by the state the brain is in when a stimulus is delivered, leading to dissociations between perception and reality. Prior research has found that the amplitude of prestimulus neural oscillations in and around the alpha frequency band (8-13 Hz) is predictive of whether a near-threshold visual stimulus or magnetic stimulation-induced phosphene is detected (Babiloni et al., 2006; Busch et al., 2009; Chaumon & Busch, 2014; Ergenoglu et al., 2004; Mathewson et al., 2009; Romei et al., 2008; Samaha, Gosseries, et al., 2017; van Dijk et al., 2008). However, changes in detection probability can come about either because an observer’s sensory system is better able to distinguish stimulus presence from stimulus absence (sensitivity change), or because an observer is more or less biased to report stimulus presence (criterion change). Recent experiments based in signal detection theory (SDT) have shown that the effect of prestimulus alpha amplitude on detection probability is explainable by a criterion change, such that observers are more likely to report a stimulus when alpha amplitude is low, even when a stimulus is absent (Iemi et al., 2017; Limbach & Corballis, 2016).

However, SDT is ambiguous with respect to how a criterion change might be related to perceptual experience. A criterion shift could come about through a change in conscious perception if, for example, during states of low alpha amplitude, the observer has a weak hallucination of a stimulus when there wasn’t one. Alternatively, criterion change may come about for post-perceptual reasons if, for instance, an observer deliberately chooses to require less evidence before deciding a stimulus was present when alpha is weak. In support of the former explanation, two recent experiments using two-choice discrimination tasks found that subjective reports of confidence (Samaha, Iemi, et al., 2017) and visibility (Benwell et al., 2017) were higher when prestimulus alpha power was low. In both studies, subjective perception effects were accompanied by null effects on discrimination sensitivity, consistent with prior SDT analyses showing no effect of alpha power on detection *d’*. Moreover, prestimulus alpha power was found not to predict a deliberate decision bias in a two-interval forced choice task, but was consistent with a bias in observer’s perceptual experience (Iemi & Busch, 2018).

Moreover, SDT is also ambiguous with respect to how a criterion change relates to underlying neural mechanisms. The same criterion effects could emerge through a change in sensory response distributions while keeping constant the absolute level of evidence needed to commit to a decision, or through a change in the absolute level of evidence, holding constant the sensory response distributions (Denison et al., 2018; Ko & Lau, 2012; Samaha et al., 2020). Growing evidence suggests that alpha amplitude biases subjective perception by acting on sensory responses, rather than on the absolute placement of criterion. For instance, high prestimulus alpha power leads to reduced early (∼70ms) visuocortical event-related potentials in humans (Iemi et al., 2019), fMRI BOLD responses in primary (Mayhew et al., 2013) and secondary (Becker et al., 2011) visual cortex, and multiunit and/or gamma-band activity in the monkey primary visual cortex (Spaak et al., 2012; van Kerkoerle et al., 2014). Thus, because alpha amplitude in primary sensory cortex is inversely related to cortical excitability, stimuli arriving during a state of low power will elicit larger sensory responses, resulting in changes to perceptual experience. However, because such fluctuations in power are spontaneous (as opposed to, for instance, attention-related), the amplification of sensory responses applies to neurons involved in representing both possible stimuli (in a 2-choice task). This account can explain changes in detection rates, confidence, and visibility, without accompanying changes in discrimination or detection sensitivity (Samaha et al., 2020)

However, prior work showing a relationship between ongoing power and subjective reports have used stimuli that preferentially engage low-level visual areas, such as luminance gratings (Samaha, Iemi, et al., 2017), or Gaussian light patches (Benwell et al., 2017). This approach is sensible given that alpha correlates negatively with low-level visual cortex excitability, but leaves open the possibility that different effects are observed when making judgments about high-level visual properties. Indeed, it has been found that alpha oscillations in macaque inferior temporal (IT) cortex (an area presumably involved in high-level perceptual representations), have a laminar profile that is distinct from V1, V2, and V4 (Bollimunta et al., 2008, 2011). Moreover, in direct opposition to early visual areas, IT alpha power shows a *positive* relationship with stimulus evoked multi-unit and gamma-band activity (Mo et al., 2011). Thus, perceptual decisions that rely on IT cortex representations may bear an opposite relationship with alpha power as compared to tasks that engage low-level visual areas.

We tested this possibility by recording electroencephalography (EEG) while observers performed a near-threshold face/house discrimination task while providing visibility judgments. Single-trial analyses of time-frequency power revealed a negative relationship between ongoing alpha-band amplitude and visibility judgments, but prestimulus activity was not predictive of decision accuracy. These results replicate and extend prior work showing that spontaneous alpha selectively impacts subjective, but not objective perceptual processing during high-level perceptual decisions. Moreover, it seems the effects of alpha on high-level perception resemble those associated with low-level perception, despite differences in the physiology of alpha in high- and low-level visual areas.

## 2. Materials and methods

### 2.1. Participants

8 observers (4 female; age range 19-27) were recruited from the University of Wisconsin-Madison community and participated for monetary compensation. Based on the effect size from our prior work (Samaha, Iemi, et al., 2017; section 3.2), this sample size achieves 80% power to detect the relationship between prestimulus alpha power and visibility. The University of Wisconsin-Madison Health Sciences Institutional Review Board approved the study protocol. All subjects provided informed consent and self-reported normal or corrected-to-normal vision and no history of neurological disorders. The EEG data reported here were collected inside an MRI scanner in the context of a simultaneous EEG/fMRI experiment.

### 2.2. Stimuli

Target stimuli were briefly presented (17 ms) grayscale images of faces and houses (Sterzer et al., 2008), which were matched with one another and with the gray background for mean luminance, and which were presented at fixation within a 2D Gaussian aperture extending 13 degrees of visual angle (DVA). We chose four unique face images and four unique house images (this small selection was necessary to estimate contrast thresholds separately for each stimulus in a timely manner; see *Procedure*). To reduce stimulus awareness, targets were backward-masked using a rapid sequence of eight full-contrast phase-scrambled versions of the face and house stimuli, lasting 200 ms (see Fig. 1A). Masking images were randomly sampled from the set of 8 textures, with the constraint that the first four and second four both contain two house textures and two face textures. We refer to this as a “dynamic mask”, which has been previously shown to be effective at masking scene gist (Loschky et al., 2010). The inter-stimulus interval (ISI) between target and mask was 33 ms. A central fixation cross was displayed throughout each trial (0.44 DVA). Stimuli were back-projected onto a screen visible from within the MRI scanner via mirrors situated above the observer’s head. The projector had a refresh rate of 60 Hz and a middle gray background (RGB vales each set to 128).

**Fig. 1.**
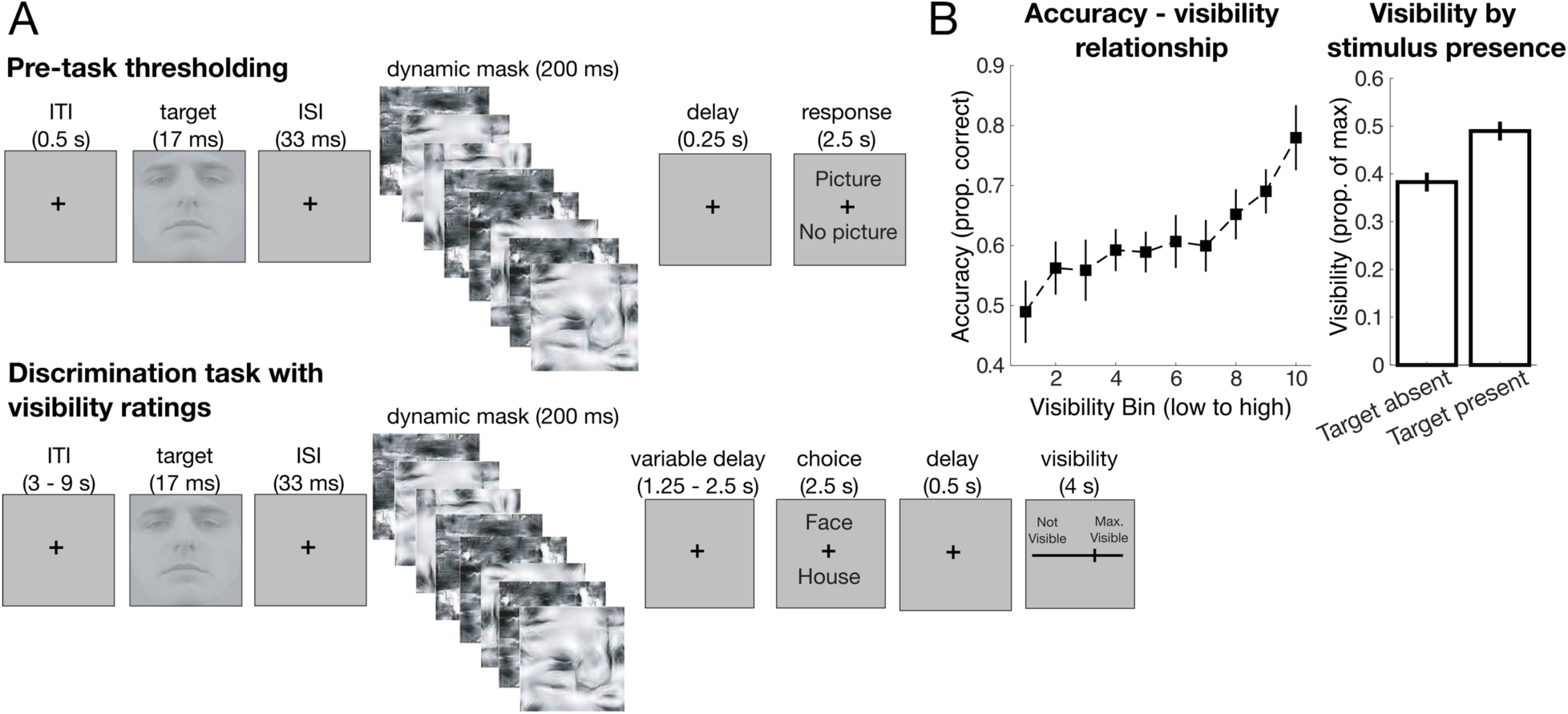
Task schematics and behavior. **A**. The pre-task thresholding procedure (top) estimated the luminance contrasts for each target stimulus that produced ∼50% subjective detections. These contrast values were then fixed throughout the main task (bottom), which consisted of a face/house discrimination with continuous visibility judgments and EEG recording. **B**. The left panel depicts mean discrimination accuracy at each of 10 visibility deciles. A clear positive monotonic relationship suggests that subjectively seen stimuli were associated with higher discrimination accuracy. The right panel shows that subjective visibility ratings were higher when an actual target stimulus was shown (80% of trials), as compared to the 20% of trials on which a target was absent. All error bars represent ±1 within-subjects SEM (Morey, 2008). ISI = inter-stimulus interval; ITI = inter-trial interval.

### 2.3. Procedure

Prior to the application of EEG, observers completed a thresholding procedure, aimed at finding target contrasts that produced approximately 50% subjectively detected targets (see Fig. 1A). Masked targets (80% of trials) were presented at varying contrast levels as determined by a one-up, one-down staircase according to observer’s detection responses (i.e., observers were asked if they perceived any image at all and did not provide discrimination judgments at this step). 20% of trials contained no stimulus. Because high-level stimuli vary along many dimensions, a 50% detection contrast-threshold was estimated for each unique face and house stimulus. The timing of experimental events is shown in Fig. 1A. Observers competed between 100 and 200 trials of this initial threshold outside of the scanner. Following the application of the EEG net, observers were situated in the MRI scanner and underwent a second round of 100 trials of the thresholding procedure, seeded with the results of the out-of-scanner threshold, to further adapt contrast levels for the display parameters within the scanner. Threshold contrasts were taken from the last reversal for each image from the in-scanner thresholding and were fixed throughout the main task.

The main task was a face/house discrimination with visibility judgments. On each trial a face (40% of trials), house (40% of trials), or no target (mask only; 20% of trials) was briefly (17ms) presented followed by an ISI (33 ms) and a dynamic mask (200 ms). After a variable delay between 1.25 and 2.5 seconds, the objective judgment was solicited when the word “face” and “house” appeared on the screen above and below fixation. Key responses were mapped to the location of each word, such that observers pressed the “up” key to select whichever choice was printed above fixation and the “down” key to indicate whichever choice was printed below fixation. The appearance of each choice option either above or below fixation was randomized on each trial to avoid a choice-selective motor preparation signal from emerging. After a 500 ms delay, subjective visibility was reported by sliding a cursor (via mouse wheel) along a visual analog scale (13 DVA) with 100 discrete points ranging from “not visible” to “maximum visibility”. The starting point of the cursor was reset to the middle of the scale on each trial. Observers were instructed to think of maximum visibility as the most visible the stimulus could be in the context of this near-threshold experiment (as opposed to the most visible *any* stimulus could ever be). After the visibility response, a fixation-only inter-trial interval (ITI) was displayed for between 3 and 9 seconds. Observers completed 280 trials of the main task, split across 7 blocks, while EEG and fMRI signals were acquired. Total experiment time was approximately 3.5 hours.

### 2.4. EEG acquisition and preprocessing

EEGs were collected at 5000 Hz from an MRI-compatible system (BrainAmp MR EEG amplifier). We used an MR-compatible electrode cap, also from Brain Vision, with 31 scalp channels plus an EKG electrode positioned on the middle of the back. Signals were referenced online to electrode AFZ. Pre-processing was done using a combination of EEGLAB13 (Delorme & Makeig, 2004) and custom MATLAB code. Several steps were then taken to clean the EEG data of MR artifacts. First, the FMRIB plug-in for EEGLAB, provided by the University of Oxford Centre for Functional MRI of the Brain (FMRIB) (Niazy et al., 2005), was used to remove artifacts caused by the MRI scanner. The gradient artifact was removed using the FASTR tool, which calculates a moving average template gradient artifact, subtracts this template from the data, then uses principal components analysis to remove up to 90% of the residual artifact left after subtraction of the template. We next downsampled the EEG data to 500 Hz in order to minimize computational demands of the next steps. The ballistocardiogram artifact was cleaned by a similar method, after a preliminary step to detect the timing of QRS complexes in the electrocardiogram. Then an initial estimation and subtraction of a template artifact was followed by reduction of residual artifact by calculation and subtraction of an optimal basis set of PCA components. A high-pass Hamming windowed-sinc FIR filter (*pop_eegfiltnew.m*) was applied at 0.1 Hz to remove slow drifts in the data and a notch-filter between 55-65 Hz removed line noise. Independent components analysis was used to remove residual MRI artifact, artifacts due to eye movements and blinks, and muscular artifact – an average of 14 components were removed from each subject’s data. After cleaning, signals were downsampled to 250 Hz, epoched from −1.2 to 1.2 seconds surrounding stimulus onset, re-referenced to the average of all channels, and baseline-corrected using the mean voltage from 200 ms prior to stimulus onset.

### 2.4. Analysis of prestimulus EEG power

Our analysis followed closely that of a prior experiment where we analyzed prestimulus power in relation to the accuracy and confidence of orientation discriminations (Samaha, Iemi, et al., 2017). We first undertook a non-parametric single-trial multiple regression analysis of time-frequency power that used single-trial accuracy and visibility judgments (normalized to each observer’s max visibility rating) to predict single-trial power across a range of prestimulus times (−500 to 0 ms) and frequencies (3 to 40 Hz). To estimate power across time and frequencies, data from each channel and trial were convolved with a family of complex Morlet wavelets spanning 3–40 Hz in 1 Hz steps with wavelet cycles increasing linearly between 3 and 8 cycles as a function of frequency. Power was obtained by squaring the absolute value of the resulting complex time series, and an analysis epoch between −500 and 0 ms relative to stimulus onset was extracted.

For each time-point, frequency, channel, and subject, beta coefficients describing the monotonic relationship between prestimulus power and accuracy (controlling for visibility) and between prestimulus power and visibility (controlling for accuracy) were estimated according the linear model:

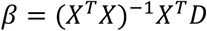

where X is the design matrix containing one column for the intercept (all 1s), one column of accuracy scores (dummy coded as 1 or 0), and one column of max-normalized visibility ratings across all trials. Superscripts *T* and −1 indicate the matrix transpose and inverse, and D is the vector of power data from all trials at a given time-frequency point and channel. Power values were first rank scored to mitigate the influence of outlying trials (this is equivalent to a Spearman correlation), and *β* was estimated using a least-squares solution. The resulting beta coefficients describe the independent contributions of accuracy and visibility to explaining prestimulus power. Beta coefficients were then converted into a z-statistic (*β*_*Z*_) relative to a subject-specific null hypothesis distribution obtained by shuffling the mapping between the data and the design matrix 2000 times and computing permuted betas (*β*_*perm*_) according to the formula described in (Cohen, 2014):

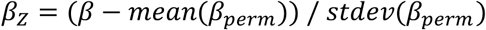

where *stdev* is the standard deviation function. *β*_*Z*_ values were then averaged over a cluster of posterior electrodes shown in Fig. 2A, which showed the highest signal amplitude in the 8-12 Hz range from between −500 and 0 ms relative to stimulus onset. These electrodes were chosen for analysis on the basis of visual inspection of group-average topography of alpha power across all trial types.

**Fig. 2.**
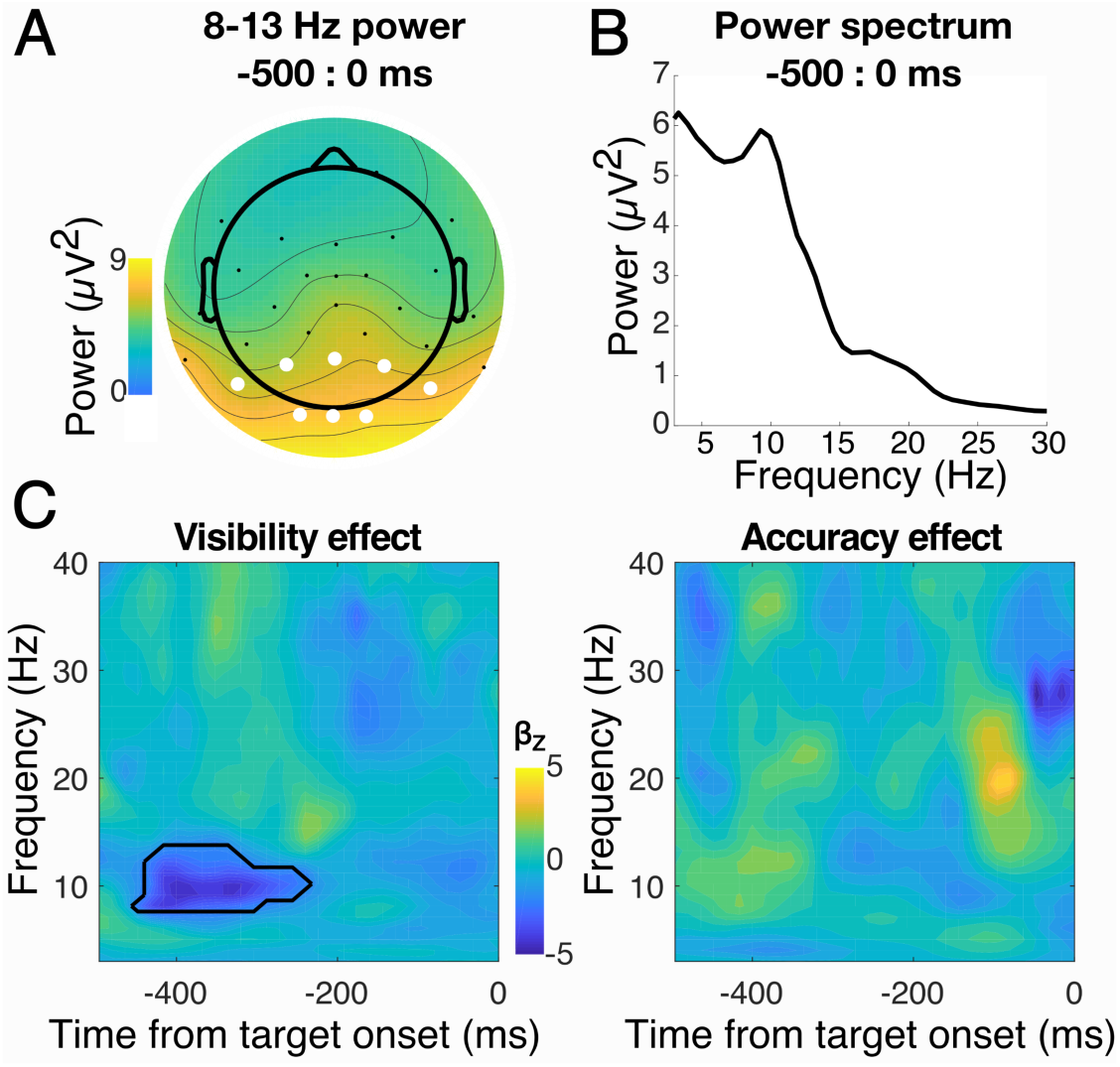
Group-averaged, single-trial multiple regression of time-frequency power. **A**. The topography of group-averaged prestimulus 8-13 Hz power revealed a posterior scalp topography. Electrodes used in subsequent analyses are shown in white. **B**. The prestimulus power spectrum of the electrode cluster shown in **A** confirms the presence of oscillatory activity with a peak at 9.3 Hz. **C**. Pseudocolor plots depicting normalized beta coefficients (*β*_*Z*_) for the effect of visibility (left) and accuracy (right) on prestimulus power across times and frequencies. Black contour lines enclose time-frequency points where *β*_*Z*_ is statistically different from zero (cluster-corrected, *α* = 0.05). A negative relationship was apparent between prestimulus alpha power and visibility, but not accuracy.

Group-level significance testing was conducted using non-parametric statistics with cluster correction to control for multiple comparisons across times and frequencies. To estimate group-level null hypothesis distributions, on each of 20,000 permutations, *β*_*Z*_ values from a random subset of subjects for both the visibility and accuracy effect were multiplied by 1 and a two-tailed *t*-test against zero was computed (this is equivalent to randomly swapping the order of the condition subtraction, e.g., A-B vs. B-A; (Maris & Oostenveld, 2007)). On each permutation, the size of the largest contiguous cluster of significant (*α* = 0.05) time-frequency points was saved, forming a distribution of cluster sizes expected under the null hypothesis. The *t*-statistics associated with the true group-level tests were converted to *z*-statistics relative to the mean and standard deviation of the empirical null hypothesis distributions and only clusters of significant *z* values (*z*_crit_ = ±1.96, *α* = 0.05) larger than the 95% percentile of the distribution of cluster sizes expected under the null hypothesis were considered significant (Cohen, 2014).

To complement our single-trial regression approach, we also binned visibility and accuracy into quartiles according to prestimulus alpha power levels obtained from a fast Fourier transform (FFT) of prestimulus data. This analysis was motivated by three considerations. (1) Time-frequency analysis necessarily results is some temporal smoothing making it is possible for poststimulus effects to smear in time and appear in the prestimulus window (Zoefel & Heil, 2013). We therefore extracted epochs between 500 and 0 ms from each trial and estimated power with a zero-padded (resolution: 0.66 Hz), Hamming window-tapered, FFT, which ensured that only prestimulus data was included in the analysis. (2) Substantial variability in peak alpha frequency exists across individuals (Haegens et al., 2014; Samaha & Postle, 2015) that is not taken into account in the time-frequency analysis. For this reason, we binned visibility and accuracy according to alpha power defined as the average over a ±2 Hz window centered on each individual’s alpha peak (identified from the power spectrum averaged across channels in our posterior region of interest). (3) Lastly, this binning analysis served as a more traditional approach by which to verify the single-trial regression results.

To statistically assess the relationship between FFT-based alpha power and visibility and accuracy, we fit linear regression slopes to each subject’s rank-scored visibility and accuracy by alpha quartiles and compared these slopes against zero at the group level with a two-tailed *t*-test. All analyses above excluded the subset of trials where no target stimulus was presented.

## 3. Results

### 3.1. Behavioral performance

We first checked whether discrimination performance and visibility judgments were sensible. As shown in Fig. 1B, face/house discrimination accuracy (proportion correct) increased monotonically with subjective visibility judgments (binned into deciles) with accuracy near chance levels (50% correct) at the lowest visibility, and up to ∼80% correct at the highest visibility. Slopes fit to individual observer’s accuracy by visibility bin data were significantly different from zero (mean slope ±SEM = 0.026±0.008; t(1,7) = 3.012, p = 0.019), indicating that accuracy increased by 2.6%, on average, for each decile increment in (max-normalized) subjective visibility. We also compared mean visibility judgments for the 20% of trials without a target stimulus to that of the remaining 80% of target trials. Visibility judgments were significantly higher for stimulus present trials (mean visibility ±SEM = 0.489±0.037), as compared to stimulus absent trials (0.383±0.051; t(1,7) = 2.689, p = 0.031), indicating that observer’s used the visibility scale appropriately.

### 3.2. Prestimulus power correlates with visibility but not accuracy

Inspection of the prestimulus power spectrum (Fig. 2B) confirmed the presence of alpha activity (peak in the power spectrum at 9.3 Hz) with a clear distribution over posterior electrodes (Fig. 2A), which we aggregated over for further analyses. Fig. 2C shows the independent contributions of visibility and accuracy in predicting time-frequency power on a single-trial basis. The pseudocolor plots show *β*_*Z*_ values averaged over observers with warm (cool) colors indicating a positive (negative) monotonic relationship between the behavioral variable and power at each time-frequency point. Contours outlined in black denote time-frequency points with significant (cluster-corrected) relationships between EEG power and behavior. As can be seen in Fig. 2C (left panel), a significant negative relationship emerged between visibility judgments and prestimulus power between 8-14 Hz and −450 and −220ms, indicating that lower prestimulus power in this time-frequency window was related to higher subsequent visibility ratings. No significant relationships were found between accuracy and prestimulus power in any time-frequency window (Fig 2C, right panel). Inspecting the time-frequency maps without cluster correction (not shown) also did not reveal a prestimulus alpha-accuracy relationship at any time point, although a transient positive relationship was apparent at 20 Hz and 100 ms prestimulus. Peak alpha frequencies ranged from 7.9 to 10.6 Hz, motivating an individually-tailored analysis of prestimulus power centered on each observer’s alpha peak. Sorting accuracy and visibility according to quartiles of FFT-based prestimulus peak alpha power (Fig. 3) revealed a negative monotonic relation between spontaneous alpha power and visibility ratings (mean slope ±SEM =-0.425±0.138; t-test against zero; t(1,7) = −3.067, p = 0.018), replicating the negative relationship found in the single-trial regression analysis. This replication suggests that the effect in the time-frequency analysis was not due to temporal smearing of post-stimulus responses because the FFT analysis considered only prestimulus data. In contrast, the monotonic relationship between prestimulus peak alpha power and accuracy was not significant (mean slope ±SEM = −1.3×10^−17^±0.192; t-test against zero; t(1,7) = −7.2×10^−17^, p = 0.999). Accuracy-power and visibility-power slopes were also significantly different from each other (t(1, 7) = 3.709, p = 0.007), indicating that prestimulus alpha power predicts visibility to a significantly greater extent than it predicts accuracy.

**Fig. 3.**
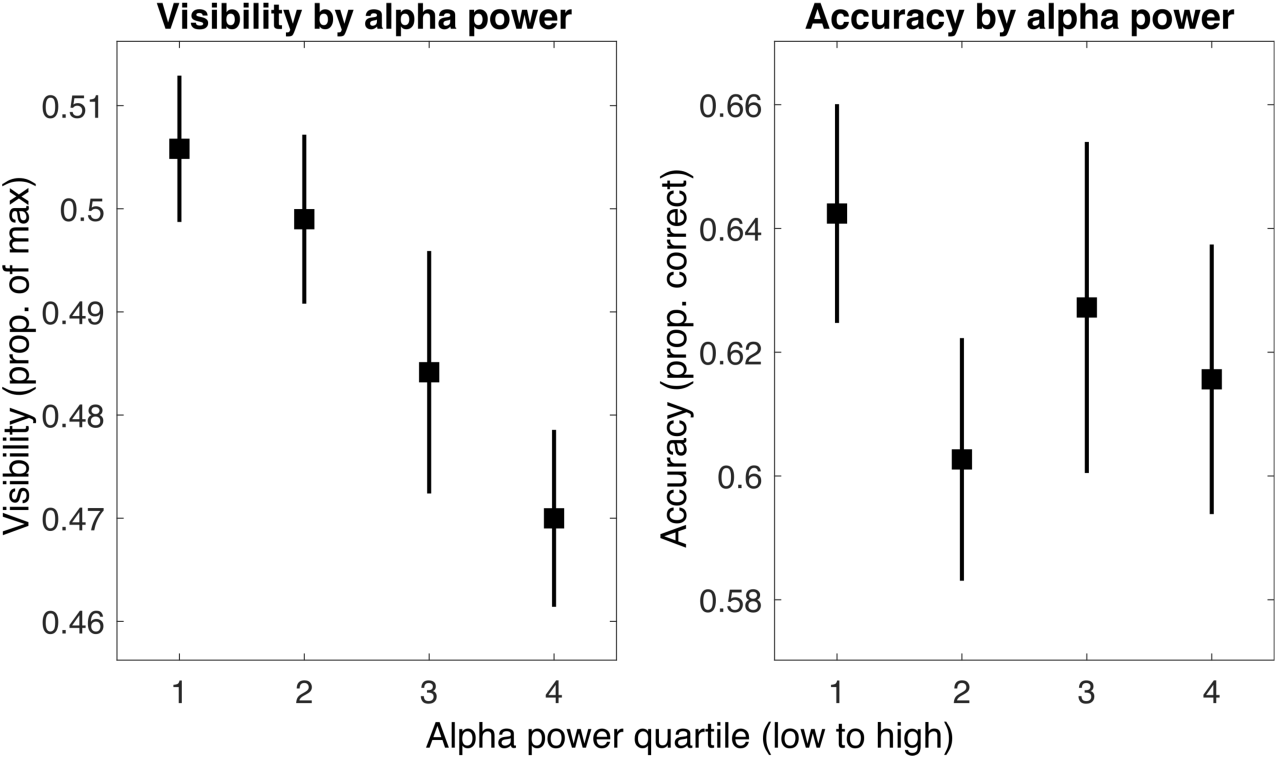
Visibility and accuracy as a function of prestimulus alpha power. FFT analysis of the prestimulus window (−500 to 0 ms) using individual peak alpha frequencies revealed the same pattern of effects: prestimulus alpha power was negatively correlated with visibility but not accuracy. Error bars indicate ±1 within-subject SEM (Morey, 2008)

## 4. Discussion

Visual perception often deviates from reality, with visual illusions and hallucinations being prime examples. Here, we show that subjective perceptual reports of complete objects are also modulated independent of stimulus factors by the amplitude of ongoing neural oscillations in the alpha range. Not only does ongoing alpha produce a dissociation between subjective perception and the stimulus, it also selectively impacts subjective visibility reports but not the veridicality of observer’s decisions. Critically, these results buttress a growing body of literature using low-level stimuli which suggests that spontaneous fluctuations in alpha-band amplitude bias the criterion for stimulus detection (Craddock et al., 2017; Iemi et al., 2017; Iemi & Busch, 2018; Limbach & Corballis, 2016) and subjective confidence/visibility (Benwell et al., 2017; Samaha, Iemi, et al., 2017; Wöstmann et al., 2019), without changing detection sensitivity (d’) or discrimination accuracy. For the first time, we show that this effect extends into a high-level perceptual discrimination task as well.

The recent “baseline sensory excitability model” (BSEM; Samaha et al., 2020) proposes that ongoing alpha power is inversely related to the excitability of early sensory brain areas and that, as a consequence, stimuli occurring during a period of weak alpha (high excitability) produce larger sensory responses. Mechanisms responsible for setting detection and visibility criteria, however, do not adapt to the increase in sensory responses and are surpassed more readily when alpha power is low. Additionally, because spontaneous fluctuations in sensory excitability as measured through EEG will, on average, equally impact populations of neurons representing both stimulus alternatives (e.g., face/house, or stimulus present/absent), the boost in sensory responses is non-specific and does not improve one’s ability to discriminate between alternative stimuli (Samaha et al., 2020). This model is premised on the observation that increased alpha power has an inhibitory influence in primary sensory cortex (Haegens et al., 2011; Iemi et al., 2019; Spaak et al., 2012; van Kerkoerle et al., 2014). In contrast, spiking activity of neurons in the high-order visual area IT in macaques have been shown to be *positively* related to alpha-band amplitude (Mo et al., 2011). Our results suggest that the inhibitory influence of alpha band activity in early sensory areas, rather than any excitatory IT influence, biases perceptual reports during a face/house discrimination task. This finding expands support for BSEM to higher-level visual perception.

Our findings provide further evidence that subjective and objective measures of visual perception are readily dissociable. The phenomenon of blindsight is a prime example of a much more profound dissociation brought about through lesions of visual cortex (Cowey & Stoerig, 1991), however our results suggest that even in neurologically intact individuals, an observer’s subjective visibility of a stimulus can be modulated independently of their ability to accurately discriminate its identity. Indeed, there is a growing body of psychophysical work pointing to the separability of subjective and objective visual reports (Koizumi et al., 2015; Lau & Passingham, 2006; Maniscalco et al., 2016; Odegaard et al., 2018; Rahnev et al., 2011; Samaha et al., 2016, 2019; Samaha & Denison, 2020; Zylberberg et al., 2012, 2014). In all of these cases (though perhaps less so in neurological blindsight), there is a question as to which metric, the objective or subjective, better tracks observers’ conscious experience. At least in the case of prestimulus alpha oscillations, the sensory origins of the bias (reviewed in Samaha et al., 2020) combined with the results of clever experimental designs aimed at ruling out decisional biases (Iemi & Busch, 2018) suggest an effect of alpha on conscious perception, per se. That is, stimuli subjectively appear more visible when alpha power is low, despite that fact this appearance does not come with more accurate decisions. If correct, dissociations such as this - between conscious perception and performance - may serve as a useful tool for further examining the neural correlates and behavioral functions of conscious perception while controlling for performance confounds (Lau & Passingham, 2006; Morales et al., 2019; Samaha, 2015)

## Acknowledgements

This work was supported by MH095984 to B.R.P

